# Synergistic effects of *APOE* ε4 and Alzheimer’s pathology on the neural correlates of episodic remembering in cognitively unimpaired older adults

**DOI:** 10.1101/2025.06.20.660774

**Authors:** Alexandra N. Trelle, Jintao Sheng, Edward N. Wilson, America Romero, Jennifer Park, Gayle K. Deutsch, Sharon J. Sha, Michael D. Greicius, Katrin I. Andreasson, Geoffrey A. Kerchner, Elizabeth C. Mormino, Anthony D. Wagner

**Author notes:** **Corresponding author:** Alexandra N. Trelle.

## Abstract

Amyloid-β (Aβ) and tau pathology begin accumulating decades before clinical symptoms and are influenced by *APOE* ε4, a key genetic risk factor for Alzheimer’s disease (AD). Although the presence of Aβ, tau, and *APOE* ε4 are thought to impact brain function, their effects on the neural correlates of episodic memory retrieval in preclinical AD remains unknown. We investigated this question in 159 cognitively unimpaired older adults (mean age, 68.9±5.8 years; 57% female) in the Stanford Aging and Memory Study. Participants completed an associative memory task concurrent with functional MRI. Aβ was measured using CSF Aβ_42_/Aβ_40_ or Florbetaben-PET imaging and tau was measured using CSF pTau_181_. Hippocampal univariate activity and cortical reinstatement – that is, reinstatement of patterns of neocortical activity that were present during memory encoding – were measured during successful memory retrieval. Analyses revealed that *APOE* ε4 was independently associated with greater Aβ and tau burden, and that associations of AD biomarkers with brain function and memory were moderated by *APOE* ε4. Among *APOE* ε4 non-carriers, Aβ burden was linked to a pattern of hippocampal hyperactivity. Among *APOE* ε4 carriers, CSF pTau_181_ was linked to weaker cortical reinstatement during memory retrieval and lower memory performance. Thus, abnormal AD biomarkers and genetic risk synergistically impact neural and behavioral expressions of memory in preclinical AD. These findings highlight the critical role of *APOE* ε4 in moderating effects of AD pathology on brain function and identify candidate mechanisms that may contribute to increased risk of memory impairment in preclinical AD.

**Significance Statement:** Hippocampus-dependent cortical reinstatement is a critical mechanism supporting episodic remembering that contributes to individual differences in memory performance in older adults. However, the contribution of early Alzheimer’s disease (AD) pathology to variability in this mechanism is unknown. We demonstrate that associations of AD biomarkers with hippocampal activity and cortical reinstatement are moderated by *APOE* ε4 in cognitively unimpaired older adults. Amyloid-β-related hyperactivity was observed in the hippocampus among *APOE* ε4 non-carriers, while CSF pTau_181_ was linked to weaker cortical reinstatement during memory retrieval and lower memory performance among *APOE* ε4 carriers. Our findings highlight synergistic effects of *APOE* and AD pathology on brain function and identify candidate mechanisms that may underlie increased risk of memory impairment in preclinical AD.

## Introduction

The neuropathological hallmarks of Alzheimer’s disease (AD), including amyloid-β (Aβ) plaques and tau neurofibrillary tangles, begin accumulating in the brain decades before clinical symptoms emerge (Braak and Braak, 1991; Jack et al., 2009). This preclinical phase of AD, characterized by abnormal Aβ biomarkers in cognitively unimpaired (CU) older adults (Price and Morris, 1999; Sperling et al., 2011; Jack et al., 2024), offers a critical window to investigate the earliest brain changes linked to AD and to develop interventions before widespread and irreversible neuronal damage occurs. Importantly, abnormal Aβ is associated with increased tau accumulation and spread (Price and Morris 1999; Knopman et al. 2021; Insel et al. 2023), which in turn is linked to greater risk of cognitive decline (Hanseeuw et al., 2019; Sperling et al., 2019; Betthauser et al., 2020; Ossenkoppele et al., 2021) and to cross-sectional memory impairments in CU older adults (Maass et al. 2018; Lowe et al. 2019; Trelle et al. 2021). Characterizing the neural mechanisms impacted by Aβ and tau in this group may help identify biomarkers that indicate risk for future decline and disease progression.

The apolipoprotein E (*APOE*) gene is the strongest genetic risk factor for sporadic AD (Corder et al., 1993; Farrer, 1997), with individuals carrying the ε4 allele exhibiting increased prevalence and earlier onset of Aβ positivity (Mishra et al., 2018; Jansen et al., 2022). Although the effects of *APOE* on Aβ production, clearance, and aggregation underlie much of this risk (Huynh et al., 2017; Yamazaki et al., 2019), growing evidence also suggests direct associations between *APOE* ε4 and tau aggregation and tau-mediated neurodegeneration (La Joie et al. 2021; Huang 2010; Shi et al. 2017; Therriault et al. 2020; Baek et al. 2020; Young et al. 2023; Koutsodendris et al. 2023). Understanding how *APOE* ε4 moderates Aβ-and tau-related effects on neural function is crucial for clarifying mechanisms of AD risk and associated memory impairment.

Prior work has largely examined the effects of AD pathology and *APOE* ε4 on task-evoked BOLD responses during memory encoding. While results have been mixed, existing evidence suggest that these factors may be linked to altered modulation of BOLD response, with observations of both hyperactivity and hypoactivity in relation to *APOE* ε4 (Bookheimer et al. 2000; Bondi et al. 2007; Filippini et al. 2009; Filippini et al. 2011; Nichols et al. 2012), Aβ burden (Mormino et al. 2012; Elman et al. 2014; Edelman et al. 2017; Sperling et al. 2009), and tau accumulation (Huijbers et al., 2019; Adams et al., 2021). These effects are often observed in the hippocampus or connected regions of the default mode network, consistent with their early vulnerability to AD pathology (Braak and Braak, 1991; Palmqvist et al., 2017). However, a key gap remains how these factors affect task-related activity and behaviorally relevant neural patterns during memory retrieval.

Cortical reinstatement describes the hippocampal-dependent reactivation of encoding-related cortical activity patterns during memory retrieval (Polyn et al., 2005; Tanaka et al., 2014). Electrophysiological evidence (Tanaka et al. 2014; Staresina et al. 2019) and human fMRI (Polyn et al. 2005; Kuhl et al. 2011; Gordon et al. 2014; Trelle et al. 2020) have demonstrated that reinstatement in the cortex is tightly linked with hippocampal activity and memory behavior, consistent with proposals that reinstated cortical patterns represent retrieved content that supports episodic remembering. While impoverished cortical reinstatement may contribute to memory deficits in aging (St-Laurent et al., 2014; Folville et al., 2020; Trelle et al., 2020), how *APOE* ε4 and early AD pathology impact this mechanism remains unknown.

The primary aim of this study was to determine whether hippocampal activity and cortical reinstatement measured using high-resolution fMRI during episodic remembering vary with *APOE* genotype and AD pathology in preclinical AD, and whether *APOE* ε4 moderates these effects. A secondary aim was to assess how genetic, molecular, and neural factors together explain individual differences in memory.

## Materials and Methods

### Participants

This study included data from 159 cognitively unimpaired (CU) older adults enrolled in the Stanford Aging and Memory Study (SAMS), a cross-sectional observational biofluid-imaging study of memory in aging and preclinical Alzheimer’s disease (e.g. Trelle et al. 2020; 2021; Sheng et al., 2025). Clinical diagnosis was determined at a clinical consensus meeting by a panel of neurologists and neuropsychologists based on a comprehensive neuropsychological battery. SAMS eligibility required a consensus diagnosis of CU and included a Clinical Dementia Rating (CDR) score^18^ of 0 and performance within 1.5 standard deviations of demographically-adjusted means of standardized neuropsychological assessments. Other eligibility criteria included normal or corrected to normal vision and hearing, right-handedness, native English speaking, and no history of neurological or psychiatric disease. All participants provided informed consent in accordance with a protocol approved by the Stanford Institutional Review Board. Participants were included in the current study if they met all eligibility criteria and had fMRI data that met all quality assurance standards (N=159). Data from an additional 10 participants were collected but not included due to excessive head motion during fMRI scanning (n=8) or fewer than five associative hit trials in the associative memory test (n=2). Data from 100 participants in the current sample were analyzed in a previous manuscript examining associations between cortical reinstatement strength, age, and episodic memory (Trelle et al., 2020).

### APOE genotyping

*APOE* genotyping was determined from whole-genome sequencing (WGS) or obtained from National Cell Repository for Alzheimer’s Disease (NCRAD) using a Fluidigm fingerprint panel. WGS was performed at the Beijing Genomics Institute (BGI) in Shenzhen, China or sequenced as part of the Stanford Extreme Phenotypes in Alzheimer’s Disease project with sequencing performed at the Uniformed Services University of the Health Sciences (USUHS) on an Illumina HiSeq platform. The Genome Analysis Toolkit (GATK) workflow Germline short variant discovery was used to map genome sequencing data to the reference genome (GRCh38) and to produce high-confidence variant calls using joint-calling. *APOE* genotype (ε2/ ε3/ ε4) was determined using allelic combinations of single nucleotide variants rs7412 and rs429358. *APOE* genotype was available in 157 participants.

### CSF Alzheimer’s disease biomarkers

CSF was collected by lumbar puncture and stored in polypropylene tubes. CSF samples were centrifuged 10 minutes at 500 x g at 22°C and aliquoted in 500 µl volumes and stored in Simport Cryo Vial 1.2 ml tubes with internal thread and O-ring (Simport Scientific, QC, Canada, Cat# T301) at -80°C until measurement. CSF AD biomarkers Aβ_42_, Aβ_40_, pTau_181_, and total tau and were measured using the fully-automated Lumipulse *G* 1200 instrument (Fujirebio US, Malvern, PA) in a single batch analysis as previously described (Trelle et al., 2021). CSF Aβ-positivity was defined using 2-cluster Gaussian mixture modelling over CSF Aβ_42_/Aβ_40_ values, yielding a cut-off of Aβ_42_/Aβ_40_ < 0.075 (Trelle et al., 2021) based on the 0.5 probability of belonging to the Aβ+ distribution. CSF data were included for 116 participants with a CSF draw within two years of their fMRI scan (n=114) or a CSF draw >2 years after fMRI if CSF Aβ_42_/Aβ_40_ was in the Aβ-distribution (n=2), the latter used only to classify participants as Aβ- and not for analysis of continuous CSF Aβ_42_/Aβ_40_ or pTau_181_.

### Amyloid PET imaging

Amyloid PET scanning using 18F-Florbetaben was completed at the Richard M. Lucas Center for Imaging at Stanford University using a PET/MRI scanner (Signa 3T, GE Healthcare). Emission data were collected between 90-110 minutes post-injection. PET images were reconstructed using TOF optimized subset expectation maximization (TOF-OSEM) with 3 iterations, 28 subsets, and 2.78 × 1.17 × 1.17 mm voxel size. Corrections were applied for detector deadtime, scatter, randoms, detector normalization, and radioisotope decay. MR attenuation correction was performed with ZTE MR imaging. PET data were reconstructed into 5-minute frames and these frames were realigned and summed. Native space FreeSurfer labels were used to extract intensity values from the co-registered summed PET data and used to create standardized uptake value ratios (SUVRs). SUVRs were created for a global cortical ROI (an average across frontal, parietal, lateral temporal, and cingulate) using a whole cerebellum reference region and converted to centiloids (Royse et al., 2021). In the absence of CSF, amyloid PET data were included for 20 participants with either a PET scan within 2 years of their fMRI scan (n=12) or a negative PET scan collected > 2 years after the fMRI scan (n=8). Amyloid PET was used only to classify participants as Aβ+ or Aβ-, defined as > 25 centiloids (Iaccarino et al., 2025; Trelle et al., 2025).

### Cognitive data

### Word-picture associative memory task

The associative memory task was administered as previously described (Trelle et al., 2020, 2021). Briefly, this task assessed memory for word–picture pairs comprising concrete nouns paired with pictures of famous faces or landmarks. The task entailed 5 alternating study and test blocks consisting of 12 word–face and 12 word–place pairs, with an additional 6 novel foils presented during test. Memory was assessed using an associative cued recall test in which participants responded to word probes with a button response, selecting “face” or “place” if they remembered the word and could recall the associated picture or picture category; “old” if they remembered the word but could not recall the associated picture or picture category; “new” if they did not remember studying the word. Associative memory performance was estimated using a sensitivity index, associative *d’*, where hits were defined as correct associative category responses to studied words and false alarms were defined as incorrect category responses to new words. Thus, associative *d’* = Z(“correct associate category” | old) − Z(“associate category” | new). A minimum of five associative hit trials was required for inclusion in the current analyses.

### Neuropsychological test delayed recall score

The delayed recall composite score was comprised of delayed recall scores from 1) the Logical Memory subtest of the Wechsler Memory Scale; 2) the Hopkins Verbal Learning Test– Revised; and 3) the Brief Visuospatial Memory Test– Revised. Composite scores were computed by first z-scoring individual subtest scores using the full SAMS sample as reference and then averaging (Trelle et al., 2020, 2021).

### MRI data acquisition and preprocessing

MR data were acquired on a 3T GE Discovery MR750 MRI scanner (GE Healthcare) using a 32-channel radiofrequency receive-only head coil (Nova Medical). Functional data were acquired using a multiband EPI sequence (acceleration factor = 3) consisting of 63 oblique axial slices parallel to the long axis of the hippocampus (TR = 2 s, TE = 30 ms, FoV = 215 mm × 215 mm, flip angle = 74°, voxel size = 1.8 × 1.8 × 2 mm). To correct for B0 field distortions, we collected two B0 field maps before every functional run, one in each phase encoding direction. Structural MRI included a whole-brain high-resolution T1-weighted anatomical volume (TR = 7.26 ms, FoV = 230 mm × 230 mm, voxel size = 0.9 × 0.9 × 0.9 mm, slices = 186).

MR data were preprocessed using fMRIPrep 23.0.0rc0 (Esteban et al., 2019) (RRID:SCR_016216), which is based on Nipype 1.8.5 (Gorgolewski et al. 2011; Gorgolewski et al. 2018) (RRID:SCR_002502). Functional images were corrected for susceptibility distortion, head motion, and slice-timing, and co-registered to the T1w structural volume. Images with motion artifacts were automatically identified as those TRs in which total displacement relative to the previous frame exceeded 0.9 mm. Trials with framewise displacement (FD) of any timepoints exceeding 0.9 mm were identified as artifacts and excluded from all analyses. Runs in which the number of artifacts identified exceeded 25% of TRs or 50% of trials, as well as runs in which FD exceeded 5 mm, were excluded. For inclusion in the current analyses, a minimum of three study and three test blocks were required for each participant. These criteria led to exclusion of eight participants with excessive head motion.

### Univariate fMRI Analysis

General linear modeling was conducted in FSL in native space on unsmoothed, retrieval phase data. Each memory outcome was modelled separately, including associative hits, associative misses, item hits, item misses, false alarms, correct rejections, and no response trials. Events were modeled as boxcar functions based on the time of stimulus onset and duration (4s) and convolved with the canonical hemodynamic response function (double gamma function). The primary contrast of interest, retrieval success effects, was defined as the difference in activity between trials on which the associate image category was successfully remembered (i.e., associative hits) and trials in which the associate image category was not remembered (i.e., associative misses, item hits, and item misses). This contrast is referred to as associative hits > misses (AH > M).

### Single-trial activity estimation

A generalized linear model (GLM) was used to estimate the activation pattern for each encoding and retrieval trial, performed in native space on unsmoothed data. A least squares single method was employed in this single-trial model, where the target trial was modeled as one explanatory variable and all other trials were modeled as another explanatory variable (Mumford et al., 2012). Each trial was modeled at its presentation time and convolved with a canonical hemodynamic response function (double gamma). This voxel-wise GLM was used to compute the activation associated with each trial. The resulting activation brain map per trial, represented as a t-statistical map (Walther et al., 2016), was then used as inputs to the classifier for multivoxel pattern analysis after being masked by each region of interest.

### Regions of interest

Univariate activity during successful associative retrieval (associative hits > misses) was extracted from the whole hippocampus, which was manually defined on T2-weighted structural scan (0.4 x 0.4 x 2mm) using established procedures (Trelle et al., 2020). Cortical reinstatement was estimated in ventral temporal cortex (VTC) and angular gyrus (ANG), two a priori regions of interest (ROIs) known to demonstrate reinstatement linked to within- and between-subject variability in memory in older and younger adults (Gordon et al., 2014; Kuhl and Chun, 2014; Favila et al., 2018; Trelle et al., 2020). ROIs were defined as described previously (Trelle et al., 2020). The VTC mask comprised three anatomical regions: parahippocampal cortex, fusiform gyrus, and inferior temporal cortex. The fusiform gyrus and inferior temporal cortex masks were generated from each participant’s Freesurfer autosegmentation volume using bilateral inferior temporal cortex and fusiform gyrus labels. These were combined with a manually defined bilateral parahippocampal cortex ROI, defined using established procedures (Trelle et al., 2020), to form the VTC mask. The ANG ROI was defined by the intersection of the Freesurfer inferior parietal lobe label and the Default Network of the Yeo 7 network atlas (Thomas Yeo et al., 2011), defined on the Freesurfer average (fsaverage) cortical surface mesh. To generate ROIs in participants’ native space from the fsaverage space label, we used the spherical registration parameters to reverse-normalize the labels and then converted the vertex coordinates of labels on the native surface into the space of each participant’s first run using the inverse of the functional to anatomical registration. Participant-specific ROIs were then defined as all voxels intersecting the midpoint between the gray-white and gray-pial boundaries. All analyses were conducted in the participants’ native space (i.e., T1 space).

### MVPA estimates of cortical reinstatement

Classification was implemented using Scikit-learn, nilearn, nibabel,, and in house Python scripts, and performed using L2-penalized logistic regression models as instantiated in the LIBLINEAR classification library (regularization parameter C =1). These models were fit to single-trial estimates, represented as a t-statistical map, from VTC and ANG. Prior to classification, the sample by voxel matrices for each region were scaled across samples within each run, such that each voxel had zero mean and unit variance. A feature selection step was also conducted, in which a participant-specific univariate contrast was used to identify the top 250 voxels that were most sensitive to each category (face, place) during encoding, yielding a set of 500 voxels over which classification analyses were performed for each region. Prior to each of 10 iterations of classifier training, the data were subsampled to ensure an equal number of face and scene trials following exclusion of trials with artefacts.

To first establish classification performance of stimulus category (face/place) during stimulus encoding, we used a leave-one-run-out n-fold cross-validation procedure on encoding phase activity patterns in each region. This yielded a value of probabilistic classifier output for each trial, representing the degree to which the encoding pattern for a trial resembled the pattern associated with a face or place trial. A continuous measure of encoding strength for each participant in each region was computed by calculating the logits (log odds) of the probabilistic classifier output and taking the mean across encoding trials.

To measure cortical reinstatement during memory retrieval, a classifier was first trained to discriminate between face vs. place encoding-phase activity patterns in each region and subsequently tested on independent retrieval-phase patterns to predict the category (face vs. place) of the remembered image on a given trial. For each ROI for each participant, a measure of reinstatement strength was derived by calculating the logits (log odds) of the probabilistic classifier output on each associative hit trial and computing mean reinstatement over trials. Reinstatement strength was signed in the direction of the correct associate for a given trial, such that, regardless of whether the trial was a face or place trial, the evidence was positive when the classifier guessed correctly, and negative when the classifier guessed incorrectly. The magnitude of reinstatement strength was thus neutral with respect to which associate category (face or place) was retrieved. To account for variance in encoding classifier performance on reinstatement, we regressed reinstatement strength (logits) on encoding classifier performance (logits) within each region, yielding a standardized residual estimate of reinstatement strength.

### Statistical analysis

All analyses were performed using R software (version 4.3.0). Linear mixed effects models including ROI (VTC, ANG), age and sex as covariates, and random intercepts to account for within-subject correlations were used to examine possible differences in associations of reinstatement strength with *APOE* ε4, Aβ status, and CSF pTau_181_ across ROIs. Given the absence of significant interactions with ROI (all *p* > 0.464; Fig. S1), a mean reinstatement score (taking the average across standardized residual reinstatement estimates in VTC and ANG) was used in the primary analyses. One outlier with a reinstatement score that was greater than 3 SD from the group mean was excluded from analyses.

Linear regression models including age and sex as covariates were used to examine cross-sectional associations of *APOE* ε4 carrier status, Aβ (status and CSF Aβ_42_/Aβ_40_) and CSF pTau_181_ with reinstatement strength and memory (i.e., associative *d’*, delayed recall memory composite). Models with memory as the outcome additionally included years of education as a covariate. Main effects of each factor were first assessed in separate models, followed by examination of interaction effects between *APOE* ε4 and Aβ (status or Aβ_42_/Aβ_40_) and CSF pTau_181_. When significant interactions were observed, models stratified by *APOE* ε4 carrier status were conducted. Participants missing *APOE* genotyping (*n*=2), Aβ status (*n*=23), or CSF Aβ_42_/Aβ_40_ and pTau_181_ (*n*=45) were excluded from models with these variables as predictors or outcomes. Due to the skewed distribution, CSF pTau_181_ concentrations were log transformed for all analyses. Statistical significance was assessed using two-sided tests with α = 0.05. Unstandardized betas and *P*-values from covariate adjusted models are reported.

## RESULTS

### APOE is associated with amyloid and pTau

A total of 159 participants enrolled in the Stanford Aging and Memory Study were included (57% females; 84% self-identified as non-Hispanic White; mean (SD) age = 68.94 (5.8) years), of which 25.6% were *APOE* ε4 carriers and 24.6% were Aβ+. Demographic characteristics for the full study sample are provided in Table 1. *APOE* ε4 carriers exhibited lower (i.e., more abnormal) CSF Aβ_42_/Aβ_40_ (β = -0.02, *p* < .001) and were more likely to be amyloid positive (β = 2.19, *p* < .0001), accounting for age and sex. *APOE* ε4 carriers also exhibited elevated CSF pTau_181_ (β = 0.21 *p =* .008), accounting for effects of age, sex, and CSF Aβ_42_/Aβ_40_ (Fig. 1).

**Table 1:** Demographic characteristics of study participants

|  | Full Sample (n=159) |
| --- | --- |
| **Age, years** | 68.9 (5.77) |
| **Sex, female** | 90 (57 %) |
| **Education, years** | 16.7 (2.01) |
| **Race** |  |
| White | 139 (87 %) |
| Asian | 12 (8 %) |
| Black or African American | 2 (1 %) |
| More Than One Race | 4 (3 %) |
| Native Hawaiian or Other Pacific Islander | 1 (1 %) |
| Unknown / Not Reported | 1 (1 %) |
| **Ethnicity** |  |
| Hispanic or Latino | 8 (5 %) |
| NOT Hispanic or Latino | 151 (95 %) |
| **APOE genotype** |  |
| $\epsilon 2/\epsilon 3$ | 12 (8 %) |
| $\epsilon 2/\epsilon 4$ | 2 (1 %) |
| $\epsilon 3/\epsilon 3$ | 105 (66 %) |
| $\epsilon 3/\epsilon 4$ | 34 (21 %) |
| $\epsilon 4/\epsilon 4$ | 4 (3 %) |
| Missing | 2 (1.3%) |
| **Amyloid status<sup>a</sup>** |  |
| Aβ- | 103 (65 %) |
| Aβ+ | 33 (21 %) |
| Missing | 23 (14.5%) |
| **Mini-mental state examination** | 29.0 (0.964) |

<sup>a</sup>Amyloid status was determined by CSF Aβ42/Aβ40 (n=116; < 0.075) or Amyloid PET (n=20; CL >25).
Values are mean (SD) for continuous variables and n (%) for categorical variables.

**Figure 1.**
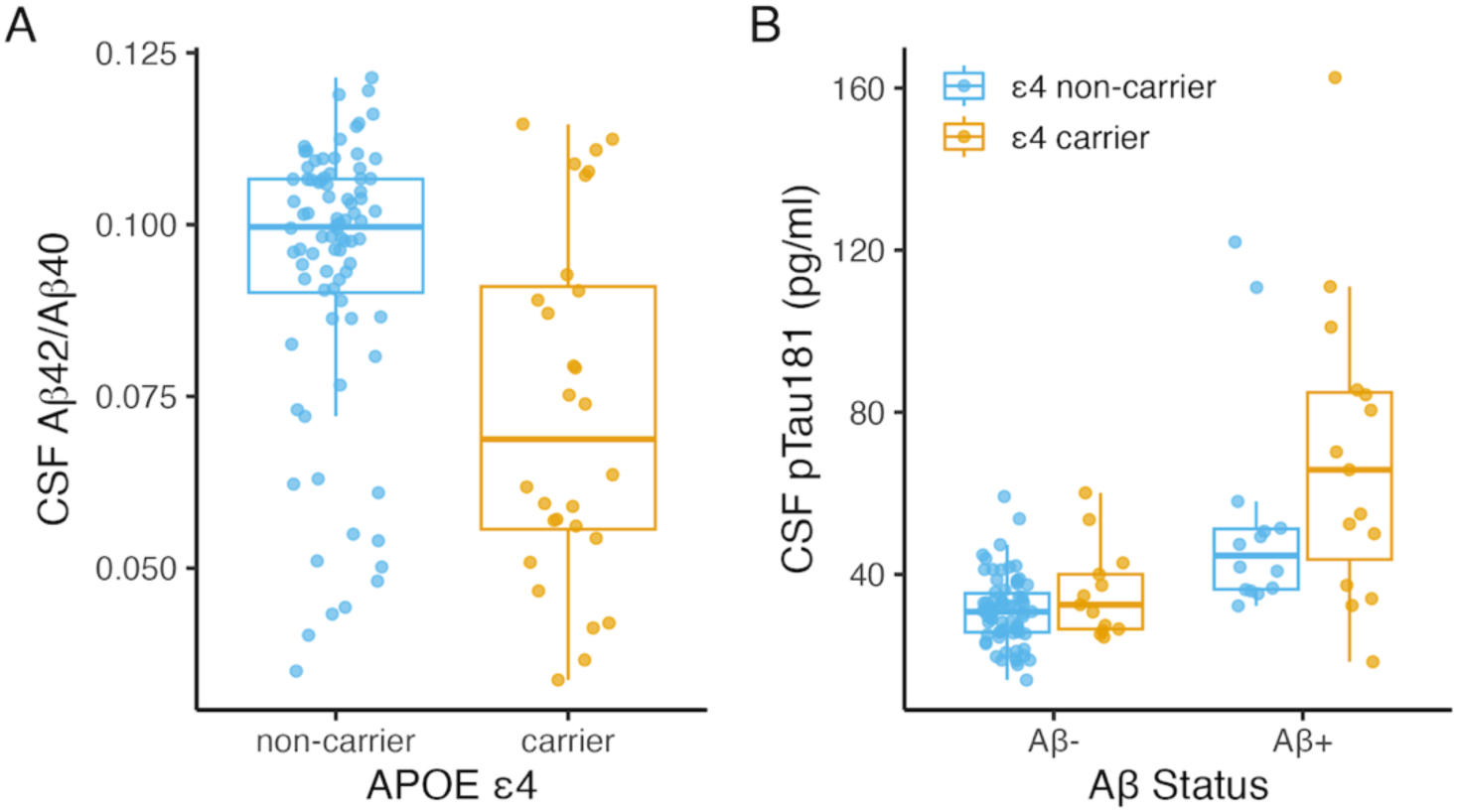
Associations between *APOE* ε4 and AD biomarkers. (**A**) Boxplots of CSF Aβ_42_/Aβ_40_ by *APOE* ε4 (*p* < .001; *n*=113) and (**B**) CSF pTau_181_ by *APOE* ε4 (*p =* .008; *n*=113), stratified by amyloid status. *P-*values were determined using linear regression models including age and sex as covariates as well as amyloid status in (**B**).

### Hippocampal activity and cortical reinstatement are associated with memory performance

We first examined associations between hippocampal activity and cortical reinstatement during successful memory retrieval and between these neural variables and memory performance, controlling for age, sex, and education. Memory was assayed with the in-scan associative memory task (Trelle et al. 2020; 2021; Sheng et al. 2025) (Fig. 2A) as well as with a memory composite score derived from a neuropsychological test battery conducted on a separate study visit (see Methods). Age was negatively associated with memory performance, estimated by associative *d’* (β= -0.30, *p* < .001; Fig. 2B) and memory composite score (β = -0.34, *p* < .001). Hippocampal univariate retrieval success effects (associative hits > all misses) did not vary with age (β = -0.004, *p* = .153; Fig. 2C), whereas cortical reinstatement was negatively related to age (β = -0.23, *p* < .005; Fig. 2D). Greater hippocampal activity was associated with stronger cortical reinstatement (β = 1.37, *p* < .001; Fig. 2E), as well as with memory performance estimated by associative *d’* (β = 1.22, *p* < .001; Fig 2F) and memory composite score (β = 0.61, *p* = .021). Cortical reinstatement strength also positively related to associative *d’* (β = 0.48, *p* < .001; Fig. 2G) and memory composite score (β = 0.23, *p* < .001). We next assessed the degree to which cortical reinstatement mediated the relationship between hippocampal activity and memory (e.g. Gordon et al. 2014; Trelle et al. 2020). In models predicting associative *d’*, both the direct (β=0.698, *p =* 0.003) and indirect (β=0.453, *p =* 0.001) effects were significant, suggesting reinstatement strength partially mediated the relationship between hippocampal activity and memory. In models predicting memory composite score, the indirect effect was significant (β=0.248, *p =* 0.015) while the direct effect was not (β=0.360, *p =* 0.173), suggesting cortical reinstatement fully mediated the relationship between hippocampal activity and memory.

**Figure 2.**
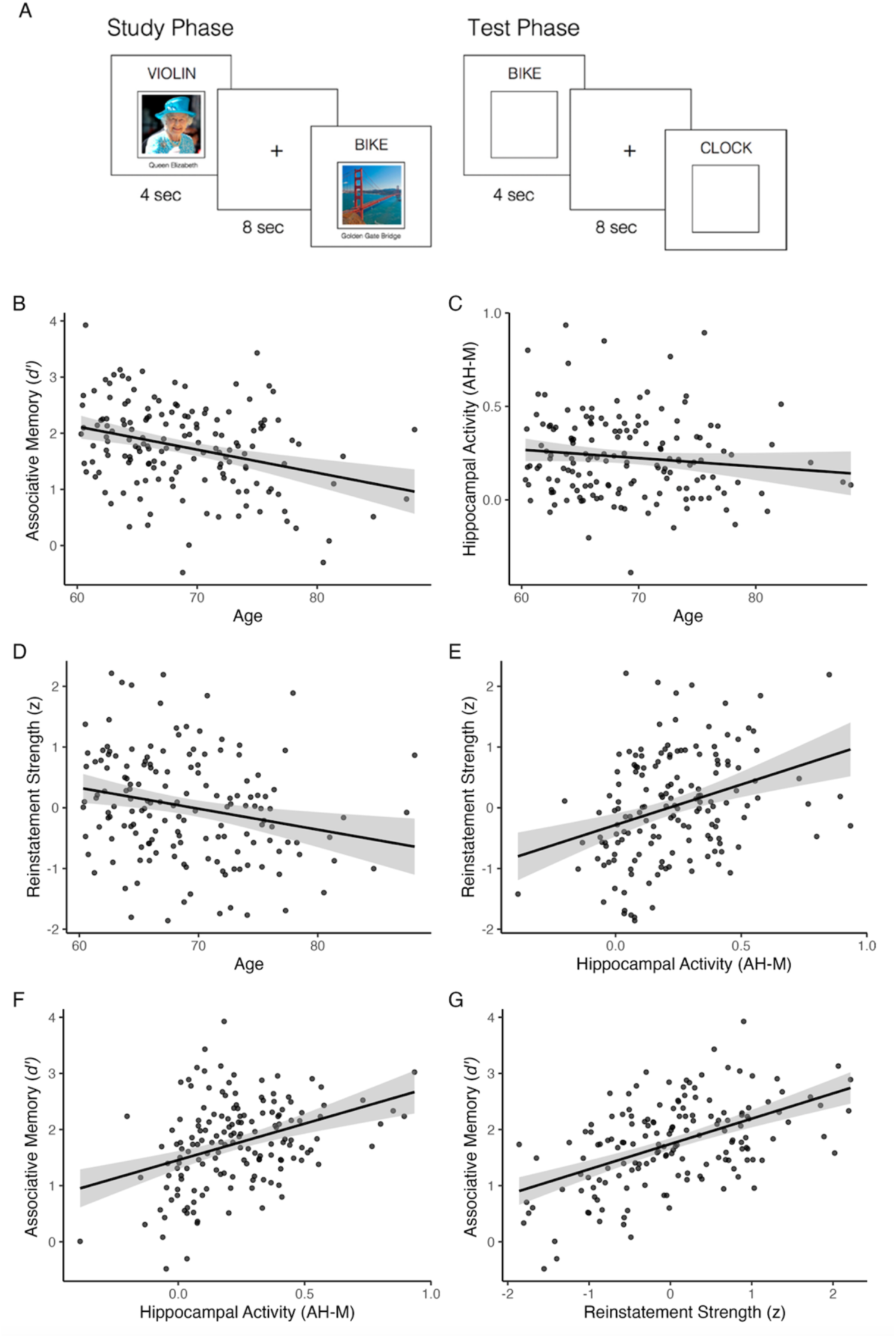
Associations between age, fMRI retrieval measures, and memory. (**A**) Task paradigm. Concurrent with fMRI scanning, participants studied word-face and word-place pairs and were tested using an associative cued-recall test in which they indicated the category associated with the presented word. (**B**) Scatterplots show associations of age with associative *d’* (*p* < .001; *n*=159), (**C**) hippocampal activity (*p* = .153; *n*=159), and (**D**) reinstatement strength (*p* < .005; *n*=158). (**E**) Scatterplots show associations of hippocampal activity with cortical reinstatement (*p* < .001) and (**F**) associative *d’* (*p* < .001) and (**G**) association of cortical reinstatement strength with to associative *d’* (*p* < .001; *n*=158). Hippocampal activity is plotted as the contrast estimate for associative hits > all misses. Reinstatement strength is plotted as the mean standardized residual, controlling for classifier accuracy during memory encoding. Plots show linear model predictions (black line) and 95% confidence intervals (shaded area). *P-*values were determined using linear regression models including age and sex as covariates, and education in models with memory as the outcome.

### Relationship between APOE ε4, AD pathology, and hippocampal activity

We next examined associations of *APOE* ε4, Aβ burden, and CSF pTau_181_ with hippocampal activity, controlling for age and sex. Main effects of *APOE* ε4, Aβ (status and continuous CSF Aβ_42_/Aβ_40_), and CSF pTau_181_ on hippocampal activity were not significant (all *p* > .322; Fig. S2). However, effects of Aβ on hippocampal activity were moderated by *APOE* ε4 (Aβ_42_/Aβ_40_ × *APOE*: *p* = .045; Aβ status × *APOE*: *p* = .061; Fig. 3AB). Among *APOE* ε4 non-carriers, there was a negative association between CSF Aβ_42_/Aβ_40_ and hippocampal activity (β = -2.35, *p* = .030), such that lower Aβ_42_/Aβ_40_ (greater amyloid burden) was linked to greater hippocampal activity. Similarly, Aβ+ ε4 non-carriers exhibited greater hippocampal activity compared to Aβ-non-carriers (β = 0.11, *p* = .047). Qualitatively, the opposite pattern was observed in *APOE* ε4 carriers, but associations with continuous Aβ_42_/Aβ_40_ (β = 1.33, *p* = .564) and Aβ status (β= -0.06, *p* = .541) were not significant. We observed trend-level evidence for moderation of CSF pTau_181_ effects by *APOE* (CSF pTau_181_ × *APOE* ε4: *p* = .055; Fig. 3C), suggesting that associations between pTau_181_ and hippocampal activity may differ in ε4 carriers, although associations were nonsignificant in each subgroup (ε4 carriers: β= -0.12, *p* = .364; non-carriers: β = 0.09, *p* = .167).

**Figure 3.**
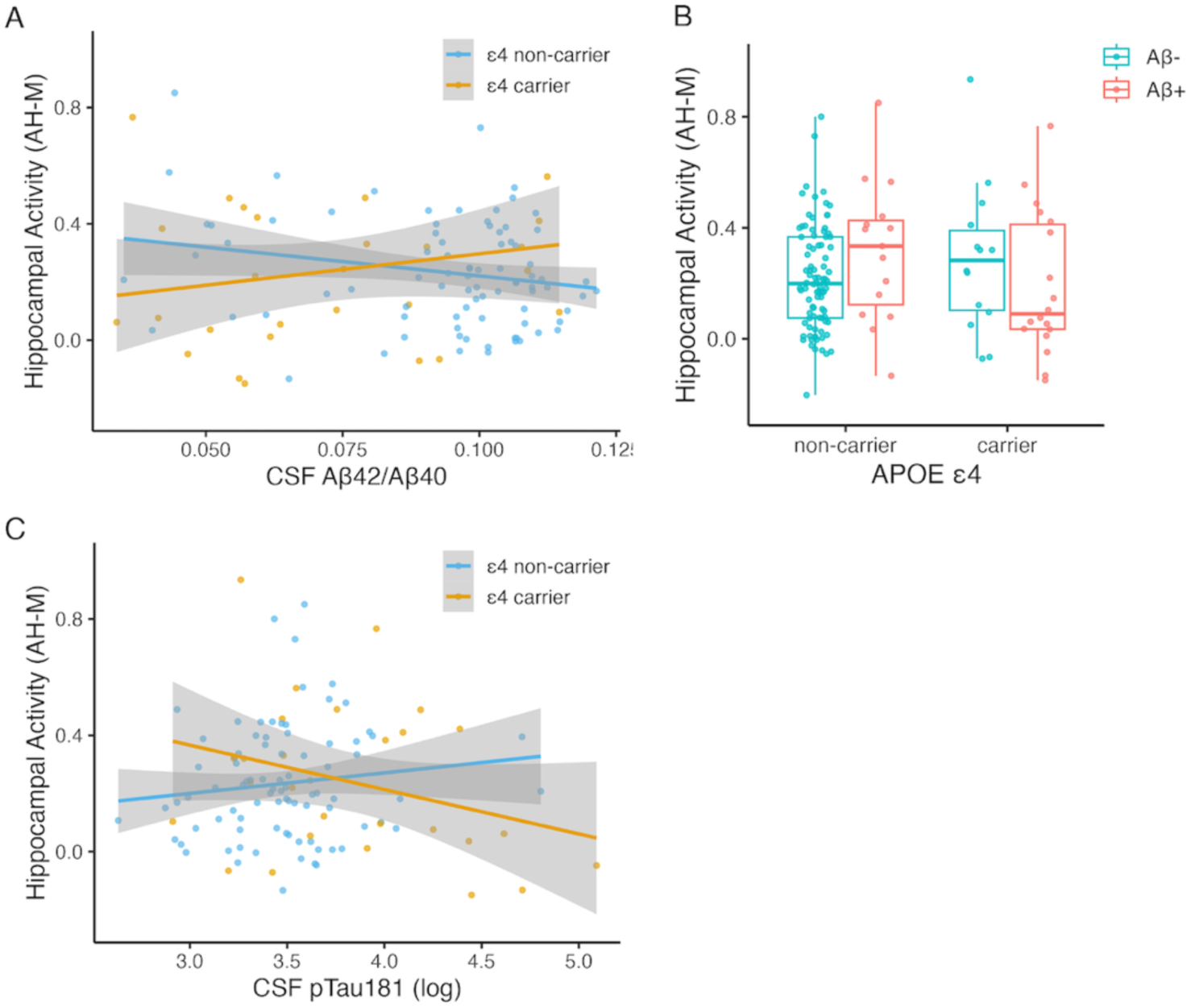
Associations of hippocampal activity with *APOE* ε4 and AD biomarkers. Interactions of *APOE* ε4 with (**A**) CSF Aβ_42_/Aβ_40_ (*p* = .045; *n*=113), (**B**) Aβ status (*p* = .061; *n*=136), and (**C**) CSF pTau_181_ (*p =* .055; *n*=113). Hippocampal activity is plotted as the contrast estimate for associative hits > all misses. Scatterplots show linear model predictions (black line) and 95% confidence intervals (shaded area). *P-*values were determined using linear regression models including age and sex as covariates.

### Relationship between APOE ε4, AD pathology, and cortical reinstatement

To further probe effects of *APOE* ε4 and AD pathology on brain function during episodic remembering, we next evaluated associations of these factors with cortical reinstatement strength, controlling for age and sex. Cortical reinstatement was weaker in *APOE* ε4 carriers (β = -0.56, *p* < .001; Fig. 4A), did not significantly vary with Aβ status (β = -0.21, *p* = .247; Fig. 4B) or CSF Aβ_42_/Aβ_40_ (β = 2.71, *p* = .468; Fig. S3A), and was negatively associated with CSF pTau_181_ (β = -0.48, *p* = .023; Fig. 4C). Associations between Aβ and reinstatement were moderated by *APOE* ε4 (Aβ × *APOE*: *p* = .027; Fig. 4D; CSF Aβ_42_/Aβ_40_ × *APOE*: *p* = .009; Fig. S3), such that effects of *APOE* ε4 on reinstatement strength were observed in Aβ+ CU (β = -1.07, *p* = .005), but not Aβ-CU (β = -0.22, *p* = .331). Similarly, the association between CSF pTau and reinstatement was moderated by *APOE* ε4 (*APOE* × pTau_181_: *p* = .007), with a significant association between pTau_181_ and reinstatement in *APOE* ε4 carriers (β = -1.08, *p* = .005) but not in non-carriers (β = 0.18, *p* = .543; Fig. 4E). The effect of pTau_181_ on reinstatement in *APOE* ε4 carriers remained significant (β = -1.15, *p* = .012) when controlling for Aβ status, which was not a significant predictor (*p* = 0.759). Moreover, when participants were stratified by *APOE* ε4, CSF pTau_181_ was the strongest predictor of reinstatement strength among ε4 carriers, explaining 26.2% of the variance, whereas hippocampal activity explained less than 1.5% of the variance (Fig. 4F). Among ε4 non-carriers, hippocampal activity was the strongest predictor of reinstatement strength, explaining 7.9% of the variance, whereas CSF pTau_181_ explained less than 1% of the variance (Fig. 4G). Taken together, these findings suggest that *APOE* ε4 moderates associations between AD biomarkers and fMRI measures of brain function during episodic remembering among CU older adults.

**Figure 4.**
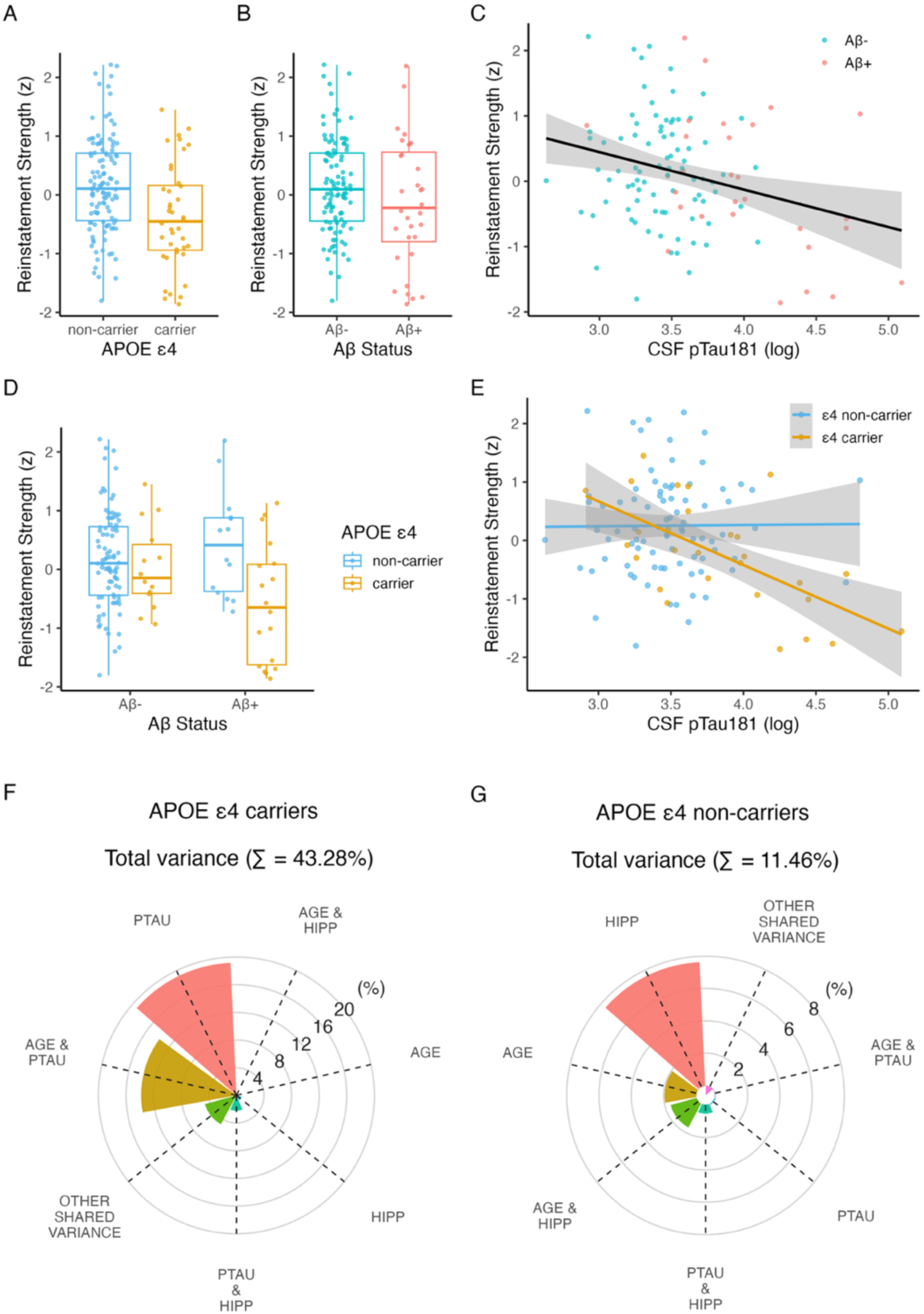
Associations of cortical reinstatement strength with *APOE* ε4 and AD biomarkers. Associations of cortical reinstatement strength with (**A**) *APOE* ε4 (*p* < .001; *n*=156), (**B**) Aβ status (*p* = .247; *n*=135), and (**C**) CSF pTau_181_ (*p* = .023; *n*=113). (**D**) Interaction of *APOE* ε4 with Aβ status (*p* = .027), such that associations with *APOE* ε4 are observed in Aβ+ (*p* = .005) but not Aβ-(*p* = .331). (**E**) Interaction of *APOE* ε4 with CSF pTau_181_ (*p* = .007), such that associations with pTau_181_ are observed in ε4 carriers (*p* = .005) but not non-carriers (*p* = .543). Reinstatement strength is plotted as the mean standardized residual, controlling for classifier accuracy during memory encoding. Scatterplots show linear model predictions and 95% confidence intervals (shaded area). *P-*values were determined using linear regression models including age and sex as covariates. (**F**) Unique and shared variance in reinstatement strength explained by age, sex, hippocampal activity, Aβ status, and CSF pTau_181_ in *APOE* ε4 carriers and (**G**) in *APOE* ε4 non-carriers.

### Relationship between APOE ε4, AD pathology, and episodic memory

Finally, we examined associations of *APOE* ε4, Aβ status, and CSF pTau_181_ with both associative *d’* (Fig. 5) and memory composite score (Fig. S4), controlling for age, sex, and education. Associative *d’* was lower in *APOE* ε4 carriers (β = -0.29, *p* = .035; Fig. 5A), marginally lower in Aβ+ individuals (β = -0.27, *p* = .070; Fig. 5B), and negatively associated with CSF pTau_181_ (β = - 0.39, *p* = .023; Fig. 5C). Effects of pTau_181_ were moderated by *APOE* (pTau_181_ × *APOE*: *p =* .002; Fig. 5E), with a negative association observed between pTau_181_ and associative *d’* in ε4 carriers (β = -1.16, *p* < .001) but not in non-carriers (β = 0.19, *p* = .429; Fig. 5E). Notably, the effect of pTau_181_ in *APOE* ε4 carriers remained significant (β = -1.27, *p* < .001) when controlling for Aβ status.

**Figure 5.**
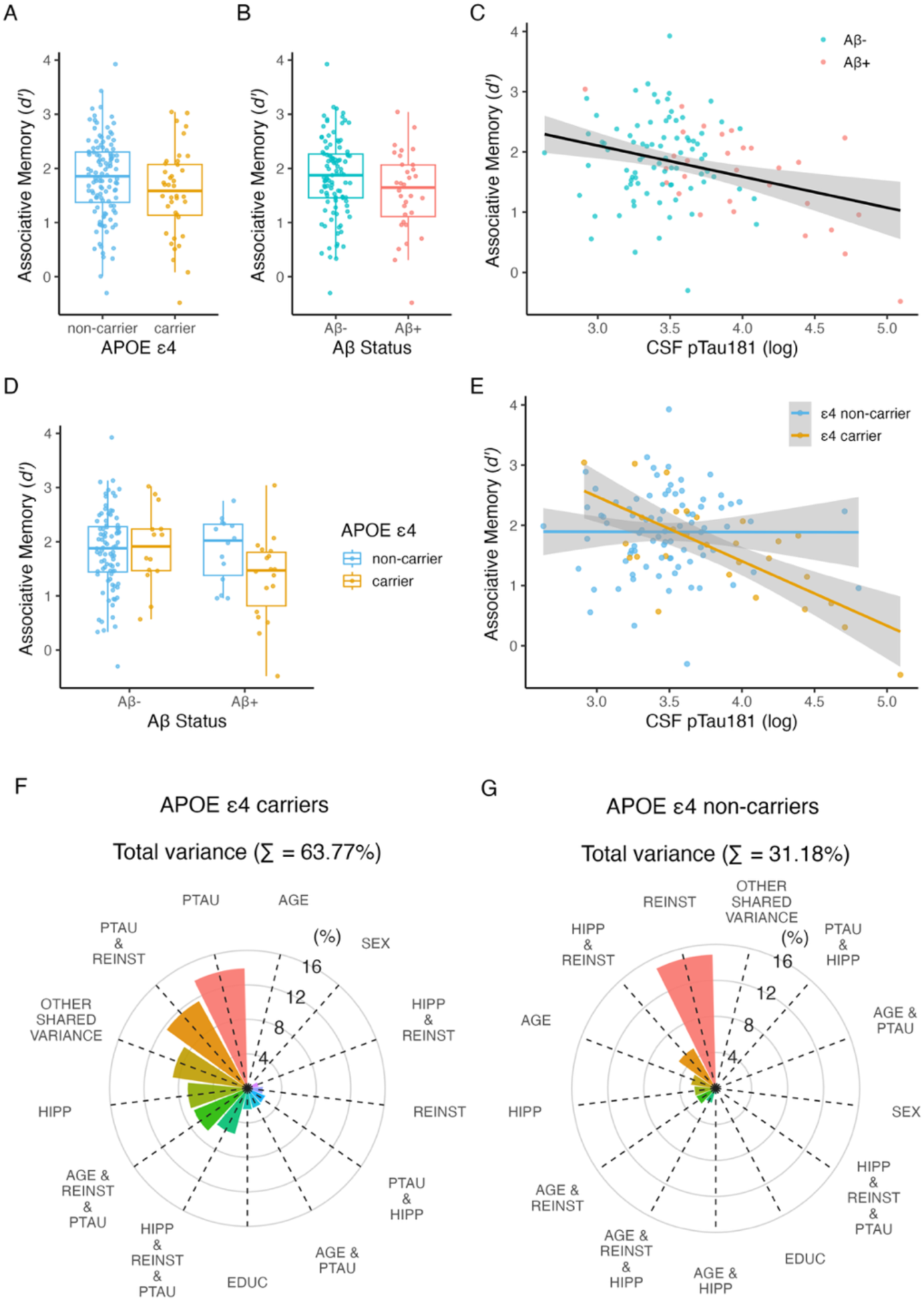
Associations of episodic memory with *APOE* ε4 and AD biomarkers. Associations of episodic memory (associative *d’*) with (**A**) *APOE* ε4 (*p* = .035; *n*=157), (**B**) Aβ status (*p* = .070; *n*=136), and (**C**) CSF pTau_181_ (*p* = .023; *n*=114). (**D**) Interaction of *APOE* ε4 with Aβ status (*p* = .224). (**E**) Interaction of *APOE* ε4 with CSF pTau_181_ (*p =* .002), such that CSF pTau_181_ was negatively related to associative *d’* in ε4 carriers (*p* < .001) but not non-carriers (*p* = .429). Scatterplots show linear model predictions and 95% confidence intervals (shaded area). *P-*values were determined using linear regression models including age, sex, and education as covariates. (**F**) Unique and shared variance in associative *d’* explained by age, sex, education, hippocampal activity, reinstatement strength, and CSF pTau_181_ in *APOE* ε4 carriers and (**G**) in *APOE* ε4 non-carriers.

When participants were stratified by *APOE* ε4 status, the factors that predicted variance in memory markedly differed between ε4 carriers and non-carriers (Fig. 5FG). Among ε4 carriers, pTau_181_ was the strongest predictor of memory, uniquely explaining 13.9% of memory variance, followed by hippocampal activity (6.9% unique variance) and reinstatement (1.8% unique variance). Moreover, consistent with the relationship between pTau and reinstatement in carriers, 11.6% memory variance was shared between these factors, and 5.5% of variance was shared between pTau, reinstatement, and hippocampal activity. In contrast, among non-carriers, pTau_181_ uniquely explained less than 1% of memory variance, hippocampal activity explained 2.4% unique variance, and reinstatement strength explained 14.9% unique variance, with 5.1% variance shared between reinstatement and hippocampal activity and less than 1% shared variance between all three factors. Collectively, these outcomes point to *APOE* ε4*-*moderated effects of pTau_181_ on memory, together with independent effects of hippocampal activity and cortical reinstatement on memory.

A qualitatively similar pattern was observed with the neuropsychological test memory composite measure (Fig. S4). Delayed recall score was lower in *APOE* ε4 carriers (β = -0.31, *p* = .020) but did not significantly vary with Aβ status (β = -0.16, *p* = .273) or CSF pTau181 (β = -0.21, *p* = .223). The association between pTau_181_ and reinstatement was again moderated by *APOE* (pTau_181_ × *APOE*: *p* = .048), reflecting a more negative association between memory and pTau181 in *APOE* ε4 carriers (β= -0.47, *p* = .287) compared to non-carriers (β= 0.27, *p* = .210), however the association was not significant in either group. This pattern is consistent with the observation that variance in the memory composite score is more weakly linked to AD pathology compared to associative *d’* (Trelle et al., 2021). Nevertheless, these results suggest that the qualitative pattern with respect to *APOE* ε4-moderated effects of pTau on memory is present across memory measures.

## DISCUSSION

The current results provide evidence for synergistic effects of *APOE* and AD pathology on neural and behavioral expressions of memory in preclinical AD. Specifically, the presence of *APOE* ε4 moderated associations of Aβ with hippocampal activity, with Aβ-related hyperactivity observed in non-carriers and the opposite patten observed in carriers. Moreover, cortical reinstatement was weaker in *APOE* ε4 carriers and individuals with elevated CSF pTau, with individuals sharing these phenotypes exhibiting the strongest negative association. Similarly, the conjunction of *APOE* ε4 and elevated pTau_181_ was associated with the greatest impairment in episodic memory across multiple assays. This work demonstrates the impact of Aβ and tau on core neural correlates of memory retrieval among CU older adults while emphasizing a key role of *APOE* in influencing effects of AD pathology on brain function and memory.

The current findings complement and extend the existing literature in two important ways. First, we build on prior observations of Aβ-associated increases in hippocampal activity during memory encoding (Sperling et al., 2009; Mormino et al., 2012; Elman et al., 2014; Edelman et al., 2017), here identifying a pattern of hyperactivity during memory retrieval. Importantly, the present data suggest that this pattern may be moderated by *APOE* ε4, with hippocampal hyperactivity scaling with greater Aβ burden only in non-carriers. This pattern is broadly compatible with findings linking both soluble and fibrillar forms of Aβ to increased neuronal excitability and aberrant neuron firing (Busche et al., 2008, 2012). Aβ-related hyperactivity was not observed among ε4 carriers, who exhibited a nonsignificant trend in the opposite direction (i.e., hypoactivity). This result may reflect observations that alterations in task-related activity as a function of *APOE* and Aβ may change across the lifespan and with disease progression. For example, prior work suggests that *APOE*-linked activity differences may vary with age, with evidence for ε4-associated hyperactivity in younger adults and hypoactivity in older adults (Filippini et al. 2009; Filippini et al. 2011). Although these prior studies were cross-sectional and did not include AD biomarkers, this pattern could reflect a transition from hyper-to hypoactivity with age and/or AD pathology. A shift from hyper-to hypoactivity also has been observed more directly in longitudinal studies (O’Brien et al., 2010), such that individuals who exhibited hyperactivity at baseline were more likely to exhibit hypoactivity and greater memory decline two years later. Thus, it is possible that older ε4 carriers are more likely to be further progressed along the AD pathological cascade (e.g., due to an earlier age of Aβ+ onset and increased tau accumulation) and thus more likely to exhibit hypoactivity compared to ε4 non-carriers. While consistent with our cross-sectional findings, a direct test of this possibility requires longitudinal data.

The present results also identify, for the first time, evidence of weakened cortical reinstatement in *APOE* ε4 carriers and individuals with elevated CSF pTau, with individuals sharing these phenotypes exhibiting the strongest association. At retrieval, cortical reinstatement reflects the hippocampal-dependent reactivation of event feature representations that were present at the time of memory encoding –– in this case features diagnostic of a retrieved image’s visual category (face vs place) – which is thought to serve as mnemonic evidence supporting memory-based decisions (Polyn et al., 2005; Gordon et al., 2014; Kuhl and Chun, 2014; Tanaka et al., 2014; Staresina et al., 2019). Consistent with this view, in a subset of the current sample we found that trial-wise strength of cortical reinstatement in ventral temporal cortex and angular gyrus is linked to hippocampal activity, probability of successful associative retrieval, and faster mnemonic decisions (Trelle et al., 2020), underscoring the key role that cortical reinstatement plays in episodic remembering. The present observation of diminished reinstatement with abnormal AD biomarkers aligns with reports of subtle memory impairment in preclinical AD, often most evident within paradigms taxing high fidelity reinstatement of event features (Webb et al., 2020; Trelle et al., 2021), and may represent one mechanism that contributes to this pattern.

The manner through which elevated tau, particularly in *APOE* ε4 carriers, disrupts cortical reinstatement remains unclear. One possibility is that the observed association with global CSF pTau_181_ captures variance related to focal tau accumulation in regions supporting event feature representations, including ventral temporal and/or inferior parietal cortex, regions that exhibit elevated tau in Aβ+ CU (Lowe et al., 2018; Insel et al., 2023). Similarly, focal tau in the medial temporal lobe could impact hippocampal pattern completion processes that drive reinstatement in the neocortex and/or pattern separation processes that support the differentiation of event representations during memory encoding. Both possibilities align with prior reports of greater tau accumulation among *APOE* ε4 carriers, particularly in the medial temporal lobe (Baek et al., 2020; Therriault et al., 2020; Young et al., 2023). Future work combining Tau PET imaging and fMRI is needed to test these and alternative hypotheses. Nevertheless, the present findings provide novel insights into mechanisms that may underlie observed associations between abnormal AD biomarkers and both poorer memory performance (Maass et al., 2018; Lowe et al., 2019; Webb et al., 2020; Trelle et al., 2021) and increased rates of memory decline among CU older adults (Hanseeuw et al., 2019; Sperling et al., 2019; Betthauser et al., 2020; Ossenkoppele et al., 2021). The observation that *APOE* ε4 moderated effects of Aβ and pTau_181_ on both neural and behavioral indices of memory function adds new evidence to a growing body of work suggesting that *APOE* ε4 exacerbates multiple pathways relevant for AD (Parhizkar and Holtzman 2022; Therriault et al. 2021; Shi et al. 2017). Work in rodent models demonstrates that *APOE* ε4 affects tau pathogenesis, neuroinflammation, and tau-mediated neurodegeneration independently of amyloid pathology (Shi et al., 2017). Consistent with such data, we found that *APOE* ε4 was associated with elevated CSF pTau_181_ even after controlling for Aβ burden. The presence of *APOE* ε4 may exacerbate inflammatory responses to degeneration and/or render neurons more susceptible to neuronal dysfunction (Parhizkar and Holtzman, 2022). Indeed, mouse models indicate that removal of *APOE* ε4 from neurons reduces tau pathology, gliosis, neurodegeneration, and neuronal hyperexcitability (Koutsodendris et al., 2023). Additional work characterizing *APOE* ε4 effects on neuroinflammation and neuronal excitability is warranted, especially in humans. Taken together, these findings underscore the importance of examining interactions between AD pathology and genetic risk when investigating cognitive and neural changes in preclinical AD.

Importantly, while the present findings reveal associations between AD genetic risk and fluid biomarkers and hippocampal activity, cortical reinstatement, and memory, they also indicate that considerable variance in these neural and behavioral expressions of memory is unrelated to the AD markers measured here, particularly among *APOE* ε4 non-carriers. It is likely that other measures of task-related neural activity (e.g., hippocampal signals at encoding; frontoparietal signals related to attention; Sheng et al., 2025), as well as other factors that change with age (e.g., synaptic density, neurotransmission, neuroinflammation, structural connectivity) will explain significant additional variance in hippocampal retrieval activity, reinstatement strength, and memory. Future work is needed to determine whether these factors covary with AD pathology or genetic risk, or whether they represent independent pathways. Nevertheless, the current results establish the impact of early AD pathology and genetic risk on two core neural indices of brain function during episodic remembering, revealing a synergistic relationship between *APOE* and AD biomarkers among CU older adults.

This study has limitations to consider. First, although the present cohort is among the largest available datasets linking task-based fMRI and AD biomarkers in a preclinical cohort, the rates of Aβ positivity and *APOE* ε4 carriage at ∼25% mean that the subsamples with AD-related phenotypes are modest. Additional replication studies that combine fMRI and AD biomarkers in large cohorts are needed. Second, the current findings are cross-sectional. Longitudinal studies are needed to determine if patterns of hyper-and hypo-activity change over time as a function of the progression of AD pathology, and whether measures of hippocampal activity and cortical reinstatement, when combined with AD biomarkers and genetic risk, improve prediction of prospective cognitive decline among CU older adults. Finally, the present cohort is predominantly non-Hispanic White and highly educated and these findings may not generalize to the broader population. Indeed, evidence suggests that the *APOE* ε4-linked risk for AD, including links between *APOE* ε4 and AD pathology, may differ by genetic ancestry (Deters et al., 2021). Thus, future work should examine interactions between genetic risk, preclinical AD, neural function, and memory in more diverse samples.

In conclusion, the current study provides novel evidence for *APOE* ε4*-*moderated associations between AD biomarkers and hippocampal activity and cortical reinstatement, core mechanisms supporting episodic memory retrieval in CU older adults. To our knowledge, this study is the first to leverage multivoxel pattern analysis to reveal an association between behaviorally relevant patterns of neural activity during memory retrieval and preclinical AD markers, highlighting a novel approach to revealing the mechanisms underlying memory impairment across the AD clinical continuum. Collectively, the present findings highlight synergistic effects of *APOE* and AD pathology on the neural correlates of episodic remembering and identify candidate mechanisms that may underlie increased risk of memory impairment in preclinical AD.

## Conflict of interest statement

The authors declare no competing financial interests.

## Acknowledgements

This work was supported by NIH K99AG075184, R01AG048076, R01AG074339, R01AG060747, and P30AG066515. The authors would like to acknowledge the support of the Stanford Center for Precision Health and Integrated Diagnostics (PHIND) and The Phil and Penny Knight Initiative for Brain Resilience.

## Notes

### Competing Interest Statement

The authors have declared no competing interest.

## References

1. Adams JN, Maass A, Berron D, Harrison TM, Baker SL, Thomas WP, Stanfill M, Jagust WJ (2021) Reduced Repetition Suppression in Aging is Driven by Tau–Related Hyperactivity in Medial Temporal Lobe. J Neurosci 41:3917–3931.

2. Baek MS, Cho H, Lee HS, Lee JH, Ryu YH, Lyoo CH (2020) Effect of APOE ε4 genotype on amyloid-β and tau accumulation in Alzheimer’s disease. Alzheimers Res Ther 12:140.

3. Betthauser TJ, Koscik RL, Jonaitis EM, Allison SL, Cody KA, Erickson CM, Rowley HA, Stone CK, Mueller KD, Clark LR, Carlsson CM, Chin NA, Asthana S, Christian BT, Johnson SC (2020) Amyloid and tau imaging biomarkers explain cognitive decline from late middle-age. Brain 143:320–335.

4. Bondi MW, Houston WS, Eyler LT, Brown GG (2007) fMRI evidence of compensatory mechanisms in older adults at genetic risk for Alzheimer disease.

5. Bookheimer SY, Strojwas MH, Cohen MS, Saunders AM, Pericak-Vance MA, Mazzio JC, Small GW (2000) Patterns of Brain Activation in People at Risk for Alzheimer’s Disease. N Engl J Med 343:450–456.

6. Braak H, Braak E (1991) Neuropathological stageing of Alzheimer-related changes. Acta Neuropathol (Berl) 82:239–259.

7. Busche MA, Chen X, Henning HA, Reichwald J, Staufenbiel M, Sakmann B, Konnerth A (2012) Critical role of soluble amyloid-β for early hippocampal Hyperactivity in a mouse model of Alzheimer’s disease. Proc Natl Acad Sci 109:8740–8745.

8. Busche MA, Eichhoff G, Adelsberger H, Abramowski D, Wiederhold K-H, Haass C, Staufenbiel M, Konnerth A, Garaschuk O (2008) Clusters of Hyperactive Neurons Near Amyloid Plaques in a Mouse Model of Alzheimer’s Disease. Science 321:1686–1689.

9. Corder EH, Saunders AM, Strittmatter WJ, Schmechel DE, Gaskell PC, Small GW, Roses AD, Haines JL, Pericak-Vance MA (1993) Gene Dose of Apolipoprotein E Type 4 Allele and the Risk of Alzheimer’s Disease in Late Onset Families. Science 261:921–923.

10. Deters KD, Mormino EC, Yu L, Lutz MW, Bennett DA, Barnes LL (2021) TOMM40-APOE haplotypes are associated with cognitive decline in non-demented Blacks. Alzheimers Dement 17:1287–1296.

11. Edelman K, Tudorascu D, Agudelo C, Snitz B, Karim H, Cohen A, Mathis C, Price J, Weissfeld L, Klunk W, Aizenstein H (2017) Amyloid-Beta Deposition is Associated with Increased Medial Temporal Lobe Activation during Memory Encoding in the Cognitively Normal Elderly. Am J Geriatr Psychiatry 25:551–560.

12. Elman JA, Oh H, Madison CM, Baker SL, Vogel JW, Marks SM, Crowley S, O’Neil JP, Jagust WJ (2014) Neural compensation in older people with brain amyloid-β deposition. Nat Neurosci 17:1316–1318.

13. Esteban O, Markiewicz CJ, Blair RW, Moodie CA, Isik AI, Erramuzpe A, Kent JD, Goncalves M, DuPre E, Snyder M, Oya H, Ghosh SS, Wright J, Durnez J, Poldrack RA, Gorgolewski KJ (2019) fMRIPrep: a robust preprocessing pipeline for functional MRI. Nat Methods 16:111–116.

14. Farrer LA (1997) Effects of Age, Sex, and Ethnicity on the Association Between Apolipoprotein E Genotype and Alzheimer Disease: A Meta-analysis. JAMA 278:1349.

15. Favila SE, Samide R, Sweigart SC, Kuhl BA (2018) Parietal Representations of Stimulus Features Are Amplified during Memory Retrieval and Flexibly Aligned with Top-Down Goals. J Neurosci 38:7809–7821.

16. Filippini N, Ebmeier KP, MacIntosh BJ, Trachtenberg AJ, Frisoni GB, Wilcock GK, Beckmann CF, Smith SM, Matthews PM, Mackay CE (2011) Differential effects of the APOE genotype on brain function across the lifespan. NeuroImage 54:602–610.

17. Filippini N, MacIntosh BJ, Hough MG, Goodwin GM, Frisoni GB, Smith SM, Matthews PM, Beckmann CF, Mackay CE (2009) Distinct patterns of brain activity in young carriers of the *APOE* -ε4 allele. Proc Natl Acad Sci 106:7209–7214.

18. Folville A, Bahri MA, Delhaye E, Salmon E, D’Argembeau A, Bastin C (2020) Age-related differences in the neural correlates of vivid remembering. NeuroImage 206:116336.

19. Gordon AM, Rissman J, Kiani R, Wagner AD (2014) Cortical Reinstatement Mediates the Relationship Between Content-Specific Encoding Activity and Subsequent Recollection Decisions. Cereb Cortex 24:3350–3364.

20. Gorgolewski K, Burns CD, Madison C, Clark D, Halchenko YO, Waskom ML, Ghosh SS (2011) Nipype: A Flexible, Lightweight and Extensible Neuroimaging Data Processing Framework in Python. Front Neuroinformatics 5 Available at: http://journal.frontiersin.org/article/10.3389/fninf.2011.00013/abstract [Accessed September 2, 2021].

21. Gorgolewski KJ, Esteban O, Markiewicz CJ, Ziegler E, Ellis DG, Notter MP, Jarecka D, Johnson H, Burns C, Manhães-Savio A (2018) Nipype. Sopware.

22. Hanseeuw BJ et al. (2019) Association of Amyloid and Tau With Cognition in Preclinical Alzheimer Disease: A Longitudinal Study. JAMA Neurol 76:915.

23. Huang Y (2010) Aβ-independent roles of apolipoprotein E4 in the pathogenesis of Alzheimer’s disease. Trends Mol Med 16:287–294.

24. Huijbers W, Schultz AP, Papp KV, LaPoint MR, Hanseeuw B, Chhatwal JP, Hedden T, Johnson KA, Sperling RA (2019) Tau Accumulation in Clinically Normal Older Adults Is Associated with Hippocampal Hyperactivity. J Neurosci 39:548–556.

25. Huynh T-PV, Davis AA, Ulrich JD, Holtzman DM (2017) Apolipoprotein E and Alzheimer’s disease: the influence of apolipoprotein E on amyloid-β and other amyloidogenic proteins. J Lipid Res 58:824–836.

26. Iaccarino L, Burnham SC, Tunali I, Wang J, Navitsky M, Arora AK, Pontecorvo MJ (2025) A practical overview of the use of amyloid-PET Centiloid values in clinical trials and research. NeuroImage Clin 46:103765.

27. Insel PS, Young CB, Aisen PS, Johnson KA, Sperling RA, Mormino EC, Donohue MC (2023) Tau positron emission tomography in preclinical Alzheimer’s disease. Brain 146:700–711.

28. Jack CR et al. (2024) Revised criteria for diagnosis and staging of Alzheimer’s disease: Alzheimer’s Association Workgroup. Alzheimers Dement 20:5143–5169.

29. Jack CR, Lowe VJ, Weigand SD, Wiste HJ, Senjem ML, Knopman DS, Shiung MM, Gunter JL, Boeve BF, Kemp BJ, Weiner M, Petersen RC (2009) Serial PIB and MRI in normal, mild cognitive impairment and Alzheimer’s disease: implications for sequence of pathological events in Alzheimer’s disease. Brain 132:1355–1365.

30. Jansen WJ et al. (2022) Prevalence Estimates of Amyloid Abnormality Across the Alzheimer Disease Clinical Spectrum. JAMA Neurol 79:228.

31. Knopman DS, Lundt ES, Therneau TM, Albertson SM, Gunter JL, Senjem ML, Schwarz CG, Mielke MM, Machulda MM, Boeve BF, Jones DT, Graff-Radford J, Vemuri P, Kantarci K, Lowe VJ, Petersen RC, Jack CR, Alzheimer’s Disease Neuroimaging Initiative (2021) Association of Initial β-Amyloid Levels With Subsequent Flortaucipir Positron Emission Tomography Changes in Persons Without Cognitive Impairment. JAMA Neurol 78:217.

32. Koutsodendris N, Blumenfeld J, Agrawal A, Traglia M, Grone B, Zilberter M, Yip O, Rao A, Nelson MR, Hao Y, Thomas R, Yoon SY, Arriola P, Huang Y (2023) Neuronal APOE4 removal protects against tau-mediated gliosis, neurodegeneration and myelin deficits. Nat Aging 3:275–296.

33. Kuhl BA, Chun MM (2014) Successful Remembering Elicits Event-Specific Activity Patterns in Lateral Parietal Cortex. J Neurosci 34:8051–8060.

34. La Joie R, Visani AV, Lesman-Segev OH, Baker SL, Edwards L, Iaccarino L, Soleimani-Meigooni DN, Mellinger T, Janabi M, Miller ZA, Perry DC, Pham J, Strom A, Gorno-Tempini ML, Rosen HJ, Miller BL, Jagust WJ, Rabinovici GD (2021) Association of *APOE4* and Clinical Variability in Alzheimer Disease With the Pattern of Tau-and Amyloid-PET. Neurology 96 Available at: https://www.neurology.org/doi/10.1212/WNL.0000000000011270 [Accessed April 5, 2024].

35. Lowe VJ et al. (2018) Widespread brain tau and its association with ageing, Braak stage and Alzheimer’s dementia. Brain 141:271–287.

36. Lowe VJ et al. (2019) Cross-sectional associations of tau-PET signal with cognition in cognitively unimpaired adults. Neurology 93 Available at: https://www.neurology.org/doi/10.1212/WNL.0000000000007728 [Accessed April 26, 2024].

37. Maass A, Lockhart SN, Harrison TM, Bell RK, Mellinger T, Swinnerton K, Baker SL, Rabinovici GD, Jagust WJ (2018) Entorhinal Tau Pathology, Episodic Memory Decline, and Neurodegeneration in Aging. J Neurosci 38:530–543.

38. Mishra S, Blazey TM, Holtzman DM, Cruchaga C, Su Y, Morris JC, Benzinger TLS, Gordon BA (2018) Longitudinal brain imaging in preclinical Alzheimer disease: impact of APOE ε4 genotype. Brain 141:1828–1839.

39. Mormino EC, Brandel MG, Madison CM, Marks S, Baker SL, Jagust WJ (2012) A Deposition in Aging Is Associated with Increases in Brain Activation during Successful Memory Encoding. Cereb Cortex 22:1813–1823.

40. Mumford JA, Turner BO, Ashby FG, Poldrack RA (2012) Deconvolving BOLD activation in event-related designs for multivoxel pattern classification analyses. NeuroImage 59:2636– 2643.

41. Nichols LM, Masdeu JC, Mattay VS, Kohn P, Emery M, Sambataro F, Kolachana B, Elvevåg B, Kippenhan S, Weinberger DR, Berman KF (2012) Interactive Effect of Apolipoprotein E Genotype and Age on Hippocampal Activation During Memory Processing in Healthy Adults. Arch Gen Psychiatry 69:804.

42. O’Brien JL, O’Keefe KM, LaViolette PS, DeLuca AN, Blacker D, Dickerson BC, Sperling RA (2010) Longitudinal fMRI in elderly reveals loss of hippocampal activation with clinical decline. Neurology 74:1969–1976.

43. Ossenkoppele R et al. (2021) Accuracy of Tau Positron Emission Tomography as a Prognostic Marker in Preclinical and Prodromal Alzheimer Disease: A Head-to-Head Comparison Against Amyloid Positron Emission Tomography and Magnetic Resonance Imaging. JAMA Neurol 78:961.

44. Palmqvist S, Schöll M, Strandberg O, Mattsson N, Stomrud E, Zetterberg H, Blennow K, Landau S, Jagust W, Hansson O (2017) Earliest accumulation of β-amyloid occurs within the default-mode network and concurrently affects brain connectivity. Nat Commun 8:1214.

45. Parhizkar S, Holtzman DM (2022) APOE mediated neuroinflammation and neurodegeneration in Alzheimer’s disease. Semin Immunol 59:101594.

46. Polyn SM, Natu VS, Cohen JD, Norman KA (2005) Category-Specific Cortical Activity Precedes Retrieval During Memory Search. Science 310:1963–1966.

47. Price JL, Morris JC (1999) Tangles and plaques in nondemented aging and preclinical Alzheimer’s disease. Ann Neurol 45:358–368.

48. Royse SK, Minhas DS, Lopresti BJ, Murphy A, Ward T, Koeppe RA, Bullich S, DeSanti S, Jagust WJ, Landau SM (2021) Validation of amyloid PET positivity thresholds in centiloids: a multisite PET study approach. Alzheimers Res Ther 13:99.

49. Sheng J, Trelle AN, Romero A, Park J, Tran TT, Sha SJ, Andreasson KI, Wilson EN, Mormino EC, Wagner AD (in press) Top-down attention and Alzheimer’s pathology impact cortical selectivity during learning, influencing episodic memory in older adults. Sci Advances.

50. Shi Y et al. (2017) ApoE4 markedly exacerbates tau-mediated neurodegeneration in a mouse model of tauopathy. Nature 549:523–527.

51. Sperling RA et al. (2011) Toward defining the preclinical stages of Alzheimer’s disease: Recommendations from the National Institute on Aging-Alzheimer’s Association workgroups on diagnostic guidelines for Alzheimer’s disease. Alzheimers Dement 7:280– 292.

52. Sperling RA et al. (2019) The impact of amyloid-beta and tau on prospective cognitive decline in older individuals. Ann Neurol 85:181–193.

53. Sperling RA, LaViolette PS, O’Keefe K, O’Brien J, Rentz DM, Pihlajamaki M, Marshall G, Hyman BT, Selkoe DJ, Hedden T, Buckner RL, Becker JA, Johnson KA (2009) Amyloid Deposition Is Associated with Impaired Default Network Function in Older Persons without Dementia. Neuron 63:178–188.

54. Staresina BP, Reber TP, Niediek J, Boström J, Elger CE, Mormann F (2019) Recollection in the human hippocampal-entorhinal cell circuitry. Nat Commun 10:1503.

55. St-Laurent M, Abdi H, Bondad A, Buchsbaum BR (2014) Memory Reactivation in Healthy Aging: Evidence of Stimulus-Specific Dedifferentiation. J Neurosci 34:4175–4186.

56. Tanaka KZ, Pevzner A, Hamidi AB, Nakazawa Y, Graham J, Wiltgen BJ (2014) Cortical Representations Are Reinstated by the Hippocampus during Memory Retrieval. Neuron 84:347–354.

57. Therriault J, Benedet AL, Pascoal TA, Mathotaarachchi S, Chamoun M, Savard M, Thomas E, Kang MS, Lussier F, Tissot C, Parsons M, Qureshi MNI, Vitali P, Massarweh G, Soucy J-P, Rej S, Saha-Chaudhuri P, Gauthier S, Rosa-Neto P (2020) Association of Apolipoprotein E ε4 With Medial Temporal Tau Independent of Amyloid-β. JAMA Neurol 77:470.

58. Therriault J, Benedet AL, Pascoal TA, Mathotaarachchi S, Savard M, Chamoun M, Thomas E, Kang MS, Lussier F, Tissot C, Soucy J-P, Massarweh G, Rej S, Saha-Chaudhuri P, Poirier J, Gauthier S, Rosa-Neto P, for the Alzheimer’s Disease Neuroimaging Initiative (2021) APOEε4 potentiates the relationship between amyloid-β and tau pathologies. Mol Psychiatry 26:5977–5988.

59. Thomas Yeo BT, Krienen FM, Sepulcre J, Sabuncu MR, Lashkari D, Hollinshead M, Roffman JL, Smoller JW, Zöllei L, Polimeni JR, Fischl B, Liu H, Buckner RL (2011) The organization of the human cerebral cortex estimated by intrinsic functional connectivity. J Neurophysiol 106:1125–1165.

60. Trelle AN et al. (2020) Hippocampal and cortical mechanisms at retrieval explain variability in episodic remembering in older adults. eLife 9:e55335.

61. Trelle AN et al. (2021) Association of CSF Biomarkers With Hippocampal-Dependent Memory in Preclinical Alzheimer Disease. Neurology 96:e1470–e1481.

62. Trelle AN et al. (2025) Plasma Aβ42 /Aβ40 is sensitive to early cerebral amyloid accumulation and predicts risk of cognitive decline across the Alzheimer’s disease spectrum. Alzheimers Dement 21:e14442.

63. Walther A, Nili H, Ejaz N, Alink A, Kriegeskorte N, Diedrichsen J (2016) Reliability of dissimilarity measures for multi-voxel pattern analysis. NeuroImage 137:188–200.

64. Webb CE, Foster CM, Horn MM, Kennedy KM, Rodrigue KM (2020) Beta-amyloid burden predicts poorer mnemonic discrimination in cognitively normal older adults. NeuroImage 221:117199.

65. Weigand AJ, Thomas KR, Bangen KJ, Eglit GML, Delano-Wood L, Gilbert PE, Brickman AM, Bondi MW, for the Alzheimer’s Disease Neuroimaging Initiative (2021) APOE interacts with tau PET to influence memory independently of amyloid PET in older adults without dementia. Alzheimers Dement 17:61–69.

66. Yamazaki Y, Zhao N, Caulfield TR, Liu C-C, Bu G (2019) Apolipoprotein E and Alzheimer disease: pathobiology and targeting strategies. Nat Rev Neurol 15:501–518.

67. Young CB, Johns E, Kennedy G, Belloy ME, Insel PS, Greicius MD, Sperling RA, Johnson KA, Poston KL, Mormino EC, for the Alzheimer’s Disease Neuroimaging Initiative, the A4 Study Team (2023) APOE effects on regional tau in preclinical Alzheimer’s disease. Mol Neurodegener 18:1.

