## Supplemental Figures 1-4 for "Synergistic effects of *APOE* ε4 and Alzheimer’s pathology on the neural correlates of episodic remembering in cognitively unimpaired older adults"

Supplementary Material

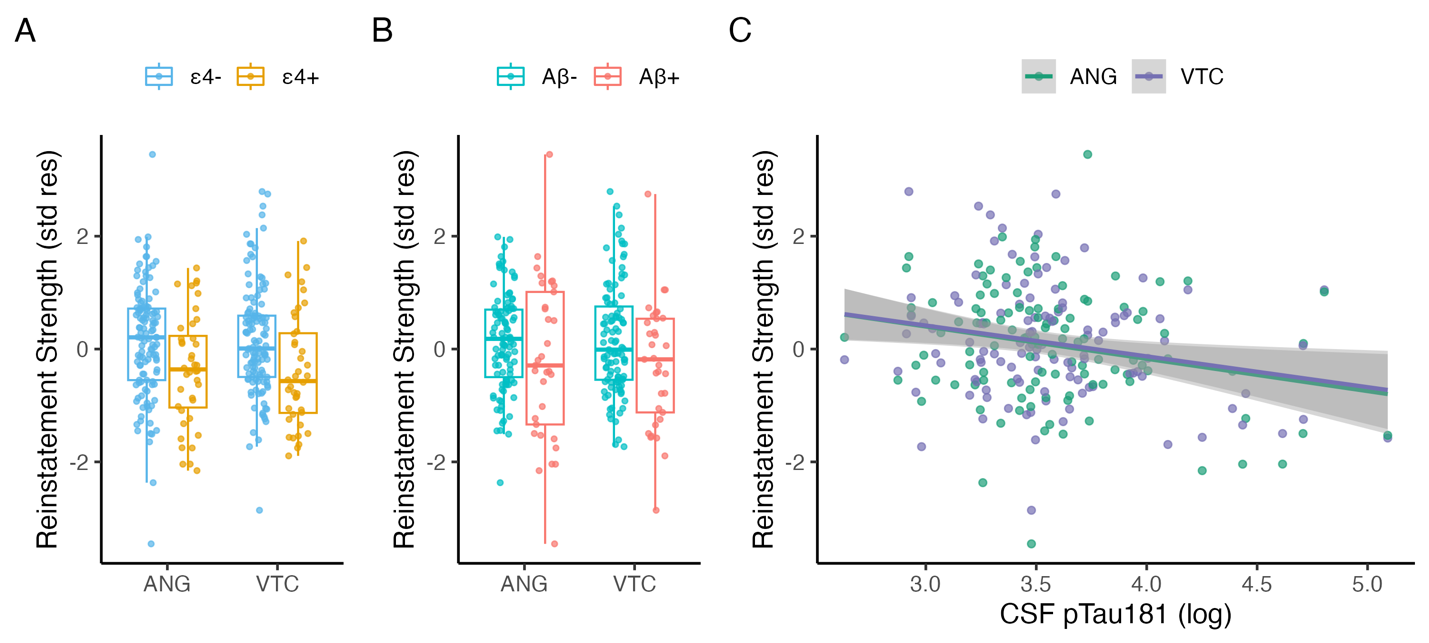

**Fig. S1 Associations of *APOE* ε4 and AD biomarkers with reinstatement by region of interest.** (**A**) Association between reinstatement strength and *APOE* ε4 (*p* < .005; *n* = 156) does not differ across regions (*p* = 0.808). (**B**) Association between reinstatement strength and Aβ status (*p =* 0.199; *n =* 135) does not differ across regions (*p =* .790). (**C**) Association between reinstatement strength and CSF pTau_181_ (*p* = 0.037; *n =* 113) does not differ across regions (*p =* 0.464). Reinstatement strength is plotted as the standardized residual controlling for classifier accuracy during memory encoding in ventral temporal cortex (VTC) and angular gyrus (ANG). Scatterplot shows linear model predictions (line) and 95% confidence intervals (shaded area). *P-*values were determined using linear regression models including age and sex as covariates.

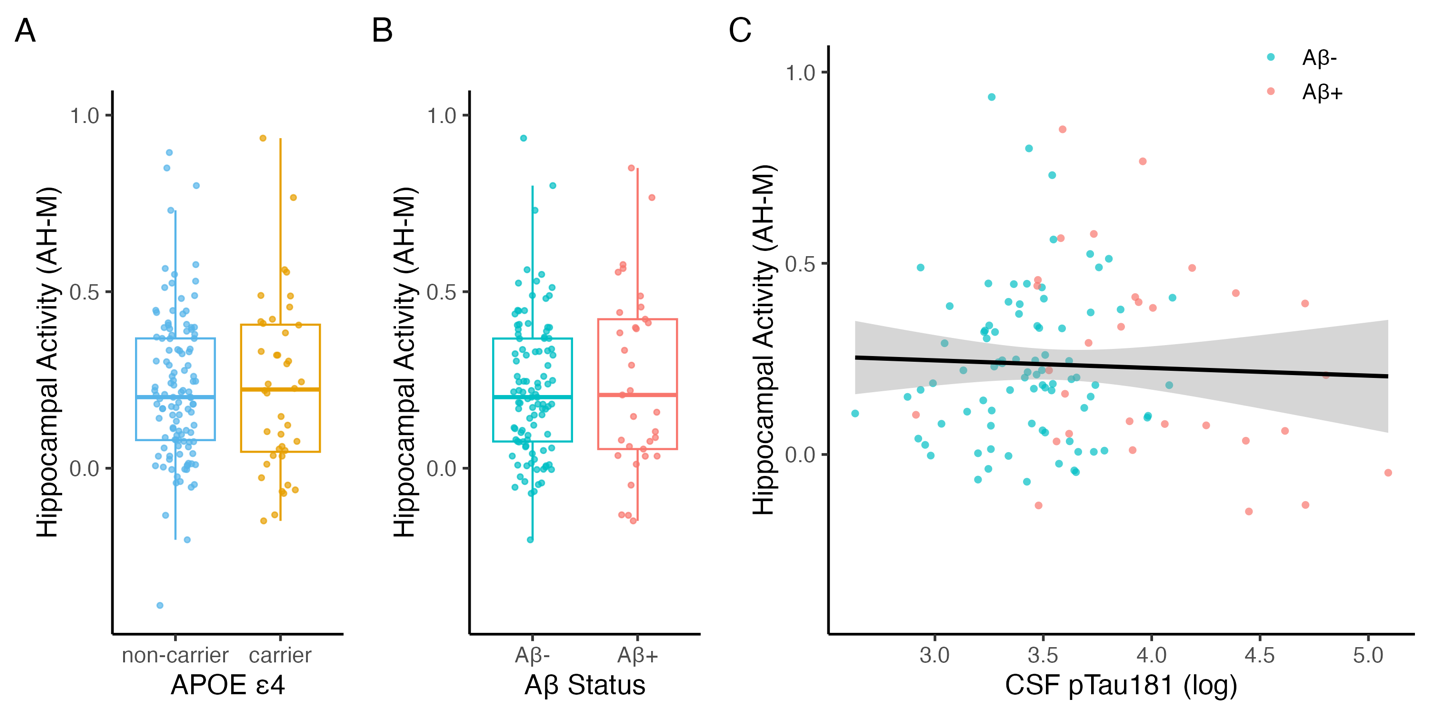

**Fig. S2 Associations of *APOE* ε4 and AD biomarkers with hippocampal activity.**

Hippocampal retrieval activity by (**A**) *APOE* ε4 (*p* = 0.995; *n =* 157), (**B**) Aβ status (*p* = 0.412; *n =* 136), or (**C**) CSF pTau181 (*p* = 0.332; *n* = 114). Hippocampal activity is plotted as the contrast estimate for associative hits > all misses. *P-*values were determined using linear regression models including age and sex as covariates.

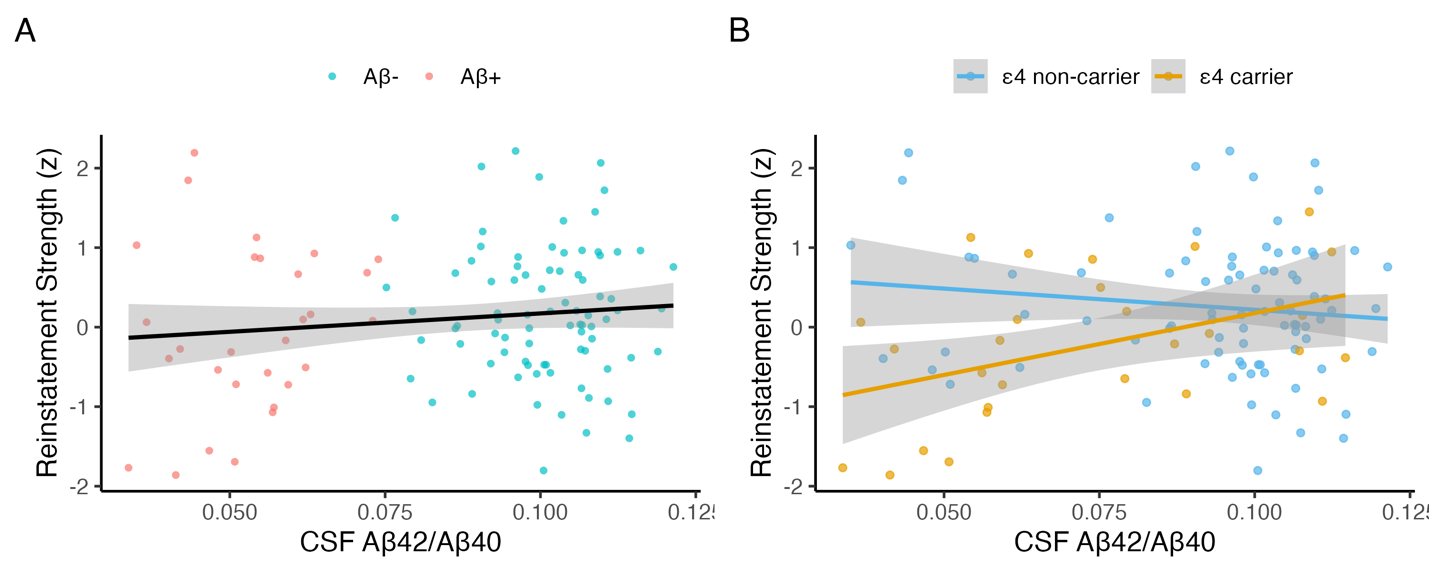

**Fig. S3 Associations of CSF Aβ_42_/Aβ_40_ with cortical reinstatement**

(**A**) Reinstatement strength by CSF Aβ_42_/Aβ_40_ (*p* = 0.468; *n* = 113). (**B**) Interaction of CSF Aβ_42_/Aβ_40_ and *APOE* ε4 on reinstatement strength (*p* = .009; *n* = 112). Reinstatement strength is plotted as the mean standardized residual, controlling for classifier accuracy during memory encoding. Scatterplot shows linear model predictions (line) and 95% confidence intervals (shaded area). *P-*values were determined using linear regression models including age and sex as covariates.

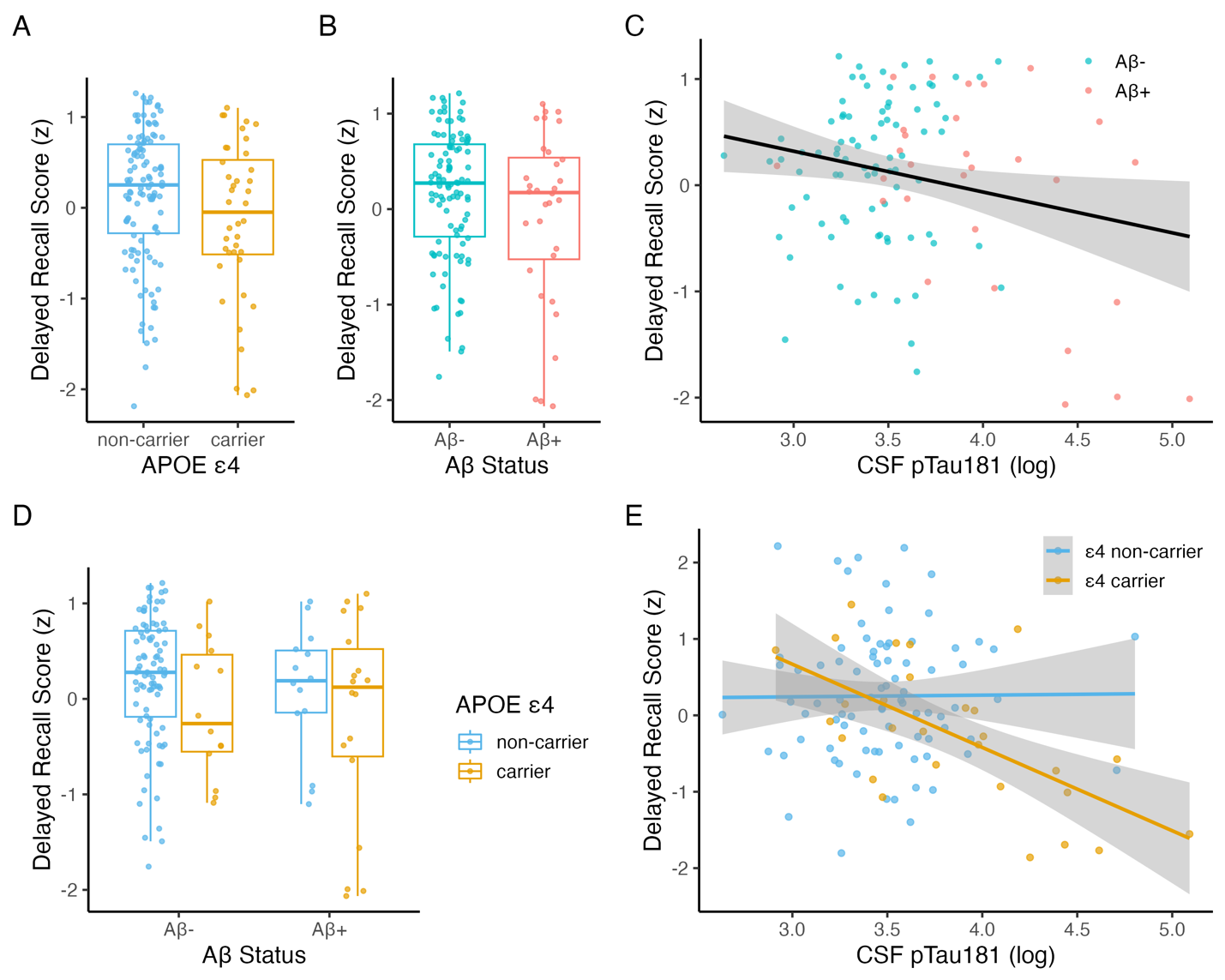

**Fig. S4 Associations of *APOE* ε4 and AD biomarkers with memory composite score.**

(**A**) Memory composite score plotted by *APOE* ε4 (*p* = .020; *n* = 157), (**B**) Aβ status (*p* = .273; *n* = 136), and (**C**) CSF pTau_181_ (*p* = .223; *n* = 114). (**D**) Interaction of *APOE* ε4 and Aβ status (*p* = .386; *n* = 135). (**E**) Interaction of *APOE* ε4 and pTau_181_ (*p* = .048; *n* = 113) indicating a stronger negative association between CSF pTau_181_ and associative memory in *APOE4* carriers relative to non-carriers. Scatterplots show linear model predictions and 95% confidence intervals (shaded area). *P-*values were determined using linear regression models including age and sex as covariates. (**F**) Unique and shared variance in associative memory explained by age, sex, education, reinstatement strength, and CSF pTau_181_ in *APOE* ε4 carriers and (**G**) in *APOE* ε4 non-carriers.
